# Vasoactive intestinal peptide confers anticipatory mucosal immunity by regulating ILC3 activity

**DOI:** 10.1101/729400

**Authors:** Cyril Seillet, Kylie Luong, Julie Tellier, Nicolas Jacquelot, Rui Dong Shen, Peter Hickey, Verena C. Wimmer, Lachlan Whitehead, Kelly Rogers, Gordon K. Smyth, Alexandra L. Garnham, Matthew Ritchie, Gabrielle T. Belz

**Affiliations:** Walter and Eliza Hall Institute of Medical Research, 1G Royal Parade, Parkville, Victoria 3052, Australia and Department of Medical Biology, University of Melbourne, Parkville, 3010, Australia; Walter and Eliza Hall Institute of Medical Research, 1G Royal Parade, Parkville, Victoria 3052, Australia and Department of Mathematics and Statistics, University of Melbourne, Parkville, Melbourne, 3010, Australia

**Keywords:** innate lymphoid cells, neuropeptide, transcriptional regulation, immune protection, neurological receptors, inflammation

## Abstract

ILC3-mediated IL-22 cytokine production is critical for the maintenance of immune homeostasis in the gastrointestinal tract. Here, we show that group 3 ILC (ILC3) constitutive function is not constant across the day but instead oscilliates between active and resting phases. Coordinate responsiveness of ILC3 in the intestine depended on food-induced expression of the neuronal hormone vasoactive intestinal peptide (VIP). Intestinal ILC3 expressed high levels of the G protein-coupled receptor, VIPR2, and activation *via* enteric neuronal VIP markedly enhanced IL-22 production and conferred gut protection. Conversely, deficiency of VIPR2 signalling led to impaired production of IL-22 by ILC3 and increased susceptibility to inflammatory gut disease. As such, intrinsic cellular rhythms synergise with the cyclic patterns of food intake to drive IL-22 thereby syncronizing intestinal epithelial protection *via* the ILC3 VIP-VIPR2 pathway.

## Introduction

The gut is the largest immune organ in the body. It harbors trillions of symbiont, commensal and pathogenic bacteria^1, 2^ that have important positive or deleterious effects on immune function, nutrient processing, and other host activities. Daily protection against the vast array of microorganisms that traverse the intestine depends on the enforced exclusion of potentially pathogenic microbes and harmful antigens derived from ingested material that could penetrate the epithelium *via* physical and immune mediated mechanisms. Rhythmic cycles within cells provide a fundamental mechanism for the body to deal with the significant fluctuations that occur in the intestine^3^. Nevertheless, protection of the intestine provides a significant challenge due to the extreme diurnal changes that occur. This necessitates coordination of physiological processes to regulate crosstalk between the microbiota and immune cells, a process essential for immune protection of the gut.

Innate lymphoid cells (ILC), particularly group 3 ILC, the most abundant ILC in the gut, are critical to neutralize invasive pathogens at the intestinal epithelial surface to ensure maintenance of homeostasis^4, 5^. ILC3-derived cytokines such as interleukin(IL)-17, IL-22 and granulocyte-macrophag colony-stimulating factor maintain immune homeostasis by triggering immune-epithelial signalling or by modulating other immune cells (e.g. monocytes) to drive tissue repair programs^4, 6, 7^. IL-22 acts mainly on non-lymphoid cells such as epithelial cells and it is essential for barrier function and tissue repair^8–10^. ILC3 are a major source of IL-22 required to protect against the inflammatory damage induced using the acute colonic model of dextran sulfate sodium (DSS)^11^, and bacterial infections such as *Clostridium difficile*^12^ and *Citrobacter rodentium*^5, 13–16^. This has led to intense efforts to understand the factors that modulate IL-22 expression for therapeutic approaches towards inflammatory intestinal diseases^17, 18^. In contrast to the positive repair functions mediated by IL-22, continuous exacerbated or uncontrolled activation of the IL-22 pathway, particularly in ILC3, leads to increased intestinal permeability^19^, chronic inflammation, an increased risk of inflammatory bowel disease^20, 21^, Crohn’s disease^22^ and colitis-associated cancer^23^. These sequelae highlight the critical requirement to tightly regulate IL-22 signalling in the mucosal epithelium. However, the mechanims that normally restrain overproduction of IL-22 in the face of continual antigenic exposure have not yet been elucidated.

Recent transcriptomic analyses have revealed that ILCs express a large array of receptors involved in the recognition of neuropeptides, hormones and metabolic signals^24, 25^. These receptors arm immune cells with the potential to act as a rheostat, integrating diverse extrinsic and intrinsic signals to finely tune immune responses^25^. For example, hormone receptors such as the androgen receptor expressed by ILC2 protect against the development of experimental asthma in male mice^26^, while the glucocorticoid receptor found on NK cells and ILC1 is triggered by lipopolysaccharide to downregulate interferon-γ production thus confering protection against septic shock^27^. A variety of other receptors including the neuroreceptors VIPR1 and VIPR2^28^, NMUR1^29–31^ and RET^11, 32^ are also expressed by ILC2 and ILC3. These receptors have been shown to be important in a tissue-specific manner in the lung and the gut highlighting the novel mechanisms utilized by ILC in integrating host responses in a complex environment. These mechanisms are especially important for ILCs as they drive their rapid secretion of effector cytokines in response to stimulation, a function central to the mobilization of early immune responses^33^. In addition, ILCs constitutively secrete mediators that are important for the homeostasis of mucosal tissues^34^ but how each of these types of receptors are regulated is not yet clear. Nevertheless, it is clear that disruption of ILCs through these pathways leads to altered immune responses with extremely harmful consequences for the organism^4^.

In the current study, we examined how innate immune cells which are strategically positioned at the intestinal interface integrate local neuronal-derived cues. We show that daily patterns of food intake coordinately regulated ILC3 responsivness. This cyclicity was only partially controlled by factors, such as circadian genes, that can intrinsically program cyclic rhythms within cells. Instead, ILC3 expression of the neuropeptide receptor vasoactive intestinal peptide receptor-2 (VIPR2) directly connected food intake with the intestinal neuronal expression of VIP to activate the protective IL-22 cytokine cascade. Collectively, we identify the mechanism governing ILC3 temporal regulation that ensures their optimal functional capacity by synergistic coordination with food intake to enforce innate defence and confer physiological protection of the gut.

## Results

### ILCs in intestinal epithelium exhibit rhythmic oscillations

Epithelial cells exhibit transcriptionally regulated diurnal fluctuations that allow temporal regulation of epithelial integrity^35, 36^. It is less clear whether ILCs, which are highly enriched in the intestinal mucosa, also exhibit cyclic patterns that connect epithelial function to immune responsiveness. To investigate whether ILCs in the gastrointestinal tract are regulated in a cyclic manner, we examined each of the ILC subsets at different stages of the light and dark cycles. All mice were fed *ad libitum* and housed under strict 24 hour dark-light conditions with lights being kept on for 12 h and time of light onset was denoted T = 0 (T0). At 4 h after the onset (T4) or offset (T16) of light, the function of the ILCs and other lymphocytes in tissues was determined (Fig. 1 and Supplementary Figs. 1-3). We first analysed the capacity of innate lymphocytes isolated at the T4 and T16 timepoints from the small intestine (Fig. 1a), the lungs and the mesenteric lymph node (LN) (Supplementary Fig. 1) to produce cytokines after stimulating the cells with PMA and ionomycin for 4.5 hours. NK cells and ILC1 did not exhibit a change in their production of interferon (IFN)-γ or tumour necrosis factor (TNF)-α among the organs tested (Fig. 1a). Distinct from this, ILC2 and ILC3 showed a difference in their capacity to produce cytokines during the day and night. IL-5-producing ILC2s were signficantly increased at T16 in both small intestine and lungs (Fig. 1a, Supplementary Fig. 1a). ILC3 showed enhanced IL-22 production specifically in the intestine but not in the mesenteric LN draining the gut, or the lungs (Fig. 1a, Supplementary Fig. 1) while IL-17 production was not markedly affected in these cells. In constrast, adaptive CD4^+^ T cells did not exhibit changes in proinflammatory cytokines such as IFN-γ, TNF-α or IL-22 while IL-17 expression showed rhymicity as has been previously observed^37^ (Supplementary Fig. 2a,b). These data suggest that among lymphocytes, ILC2 and ILC3 functions are particularly sensitive to temporal regulation.

**Fig. 1.**
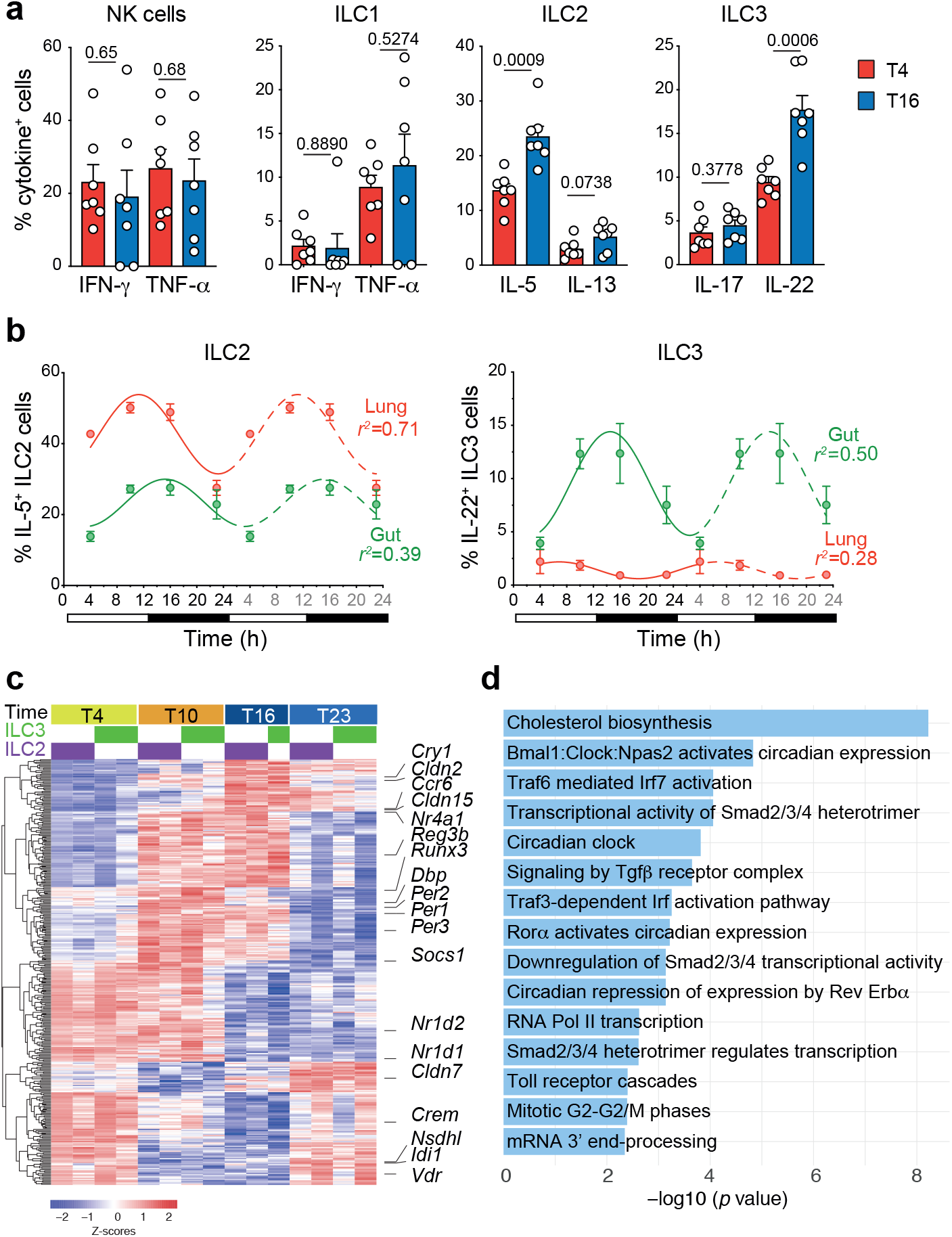
ILC2 and ILC3 activity oscilliate during the active and resting phase at steady-state in wild-type mice. **a**, Frequency of cytokine producing ILC subsets isolated from the small intestine at T4 and T16 after 4.5h of stimulation with PMA and ionomycin. Individual responses together with mean ± s.e.m. are shown (n = 7 mice at each time point). One experiment of five independent experiments with similar results is shown. **b**, Frequency of IL-5 and IL-22 constitutively produced by ILC2 (left panel) and ILC3 (right panel) from the lung and the small intestine at T4, T10, T16 and T23. The dotted line projects duplicated data to better appreciate the circadian pattern. Data show the mean and s.e.m. (*n* = 4-6 mice at each time point) for one of two independent experiments. **c,d** RNA-seq analyses of ILC2 and ILC3 in the small intestine. **c**, Heat map depicts the common genes that follow a circadian pattern in combined ILC2 and ILC3 which were identified by expression of a periodic signal using a single harmonic sinusoidal model. Individual biological replicates are shown. **d**, Top 15 pathways that oscilliate identified in ILC2 and ILC3 using Reactome analyses (https://reactome.org/).

To better understand the nature of the variation in IL-5 and IL-22 production, we analysed the constitutive cytokine expression of the ILC2 and ILC3 at four different time points during the day (Fig. 1b). We found that production of IL-5 and IL-22 by ILC2 and ILC3, respectively, followed a diurnal pattern that peaked between T14 and T16. This indicates that the individual effector programs in ILC2 and ILC3 are regulated in a time- and tissue-specific manner. As IL-5 secretion by ILC2 has previously been reported to vary during the day^28^, we focused our analysis on IL-22 secretion by ILC3. In order to elucidate the mechanisms that might characterise the selective rhythmic expression of cytokines in intestinal ILCs, we performed RNA-seq analyses on flow cytometrically-purified ILC2 and ILC3 at T4, 10, 16 and 23 over a 24 h period from the small intestine (Fig. 1c). We combined both data sets to elucidate the core set of genes that oscilliated in ILC2 and ILC3. Rhythmic changes in gene expression across the four timepoints were identified using a single harmonic sinusoidal model to fit log2 transformed data to characterise the expression of a periodic pattern amongst ILC transcripts (FDR adjusted *p*-value < 0.05). Amongst ILC2 and ILC3 this showed enrichment of genes involved in regulation of the circadian rhythm (e.g. *Dpb*, *Per1*,*2 and 3*, *Nr1d1*, *Nr1d2*, *Cry1*) indicating that ILC2 and ILC3 possess functional circadian molecular machinery. Three of the the top 15 pathways identified using Reactome analyses (https://reactome.org/) were related to circadian rhythm (Fig. 1d). Interestingly, genes involved in intestinal tissue integrity such as the claudin family (*Cldn2*, *Cldn15* and *Cldn7*) and antimicrobial proteins *Reg3b* also exhibited oscillation and were upregulated during the night time (Fig. 1c). Reactome analysis revealed that TGF-β signalling and SMAD transcriptional activity (*Socs1*, *Ncor2*, *Trim33*) were upregulated during the day. This increase in TGF-β signalling during the day could indicate an increase of the anti-infllammatory properties of the ILCs during the resting phase of the rodent. Finally, genes involved in metabolism, in particular enzymes involed in the cholesterol biosynthesis (*Msmo1*, *Mvk*, *Nsdhl*, *Idi1*, *Fdps Dhcr24*, *Sqle*) were also increased in the active phase. Collectively, this analysis revealed that ILCs exhibit significant cyclicity in gene regulation especially amongst genes that are central to mucosal immune protection. It also revealed unexpectedly that distinct functions of ILCs are regulated across the day in a time-dependent manner to favour their pro or anti-inflammatory properties.

### Intrinsic clock genes only partially regulate rhythmic ILC cytokine responses

To understand how this rhythmicity in ILC function is regulated we examined the contribution of the circadian clock gene *Bmal1* (also known as *Arntl* or Aryl hydrocarbon receptor nuclear translocator-like protein-1) that plays a key role in intrinsically regulating cellular oscillations, particularly in cytokine production. BMAL1 rhythmically binds to DNA to activate the expression of the core clock genes Period (PER), Cryptochrome (CRY1), REV-ERBα and ROR^3^. We crossed the *Il7rCre* × *Bmal1*^*flox/flox*^ conditional mice to generate the *Bmal1*^*ΔIL7R*^ strain in which *Bmal1* was deleted in all lymphoid lineages (data not shown). Mixed bone marrow chimeric mice were generated by transplanting a 1:1 mixture of wild-type (CD45.1^+^CD45.2^+^, F1) and BMAL1-lymphoid deficient bone marrow (CD45.2^+/+^*Bmal1*^*ΔIL7R*^) into lethally irradiated CD45.1^+/+^ wild-type recipients (Fig. 2a). This allowed us to simultaneously examine the responsiveness of *Bmal1*-sufficient and replete ILC subsets derived from different donors but that were exposed to identical cellular environments during tissue reconstitution. ILC subset reconstitution, cytokine production and expression patterns were examined in mice after 8 weeks (Fig. 2a-c). Both wild-type and *Bmal1*^*ΔIL7R*^ bone marrow were able to reconstitute adaptive (Fig. 2a) and most innate lymphoid subsets in the small intestine. Notably, the frequency of Rorγt^+^ ILC3 were significantly diminished in the *Bmal1*^*ΔIL7R*^ compartment (Fig. 2a,b). A significant reduction was also observed in the ability of *Bmal1*^*ΔIL7R*^ ILC3 to constitutively produce IL-22, or in response to IL-7 and IL-23, compared with wild-type ILC3 (Fig. 2c). To test whether this defect was intrinsic to the ILC3, we purified ILC3 from wild-type and the *Bmal1*^*ΔIL7R*^ mice (Supplementary Fig. 4) and analysed the production of IL-22 after 16 hours incubation with IL-7 (constitutive cytokine production), or a combination IL-7 and IL-23 to stimulate cytokine production (Fig. 2d). In both conditions, ILC3 purified from *Bmal1*^*ΔIL7R*^ mice produced less IL-22 compared with wild-type ILC3. Thus, BMAL1 is required for optimal secretion of IL-22, however, whether BMAL1 generally controls the cyclic expression of cytokines in ILC was not clear. To understand this, we analysed IL-5 production in ILC2 isolated from the small intestine of wild-type and *Bmal1*^*ΔIL7R*^ mice at T4 and T16 (Fig. 2e,f). Temporal analyses of cytokine production revealed that loss of BMAL1 did not substantially influence the overall production or the oscillatory expression pattern of IL-5 by ILC2 (Fig. 2e, left panel). This was in contrast to *Bmal1*^*ΔIL7R*^ ILC3 where we observed a marked reduction in constitutive secretion of IL-22 and a significantly diminished amplitude of the oscillation wave although the capacity to produce cytokine was not completely abolished (Fig. 2f, left panel). The provision of IL-7 and IL-23 *in vitro* increased the responsiveness of both WT and *Bmal1*-deficient ILC2 by ~ 1.3-fold (Fig. 2e, right panel), while the frequency of ILC3 able to produce IL-22 was increased ~6.5-fold. Despite the fact that stimulation with IL-23 did not restore IL-22 production in *Bmal1*-deficient ILC3 to wild-type levels (Fig. 2f, right panel), the oscilliation in cytokine production remained although the amplitude of cyclic production was reduced. In contrast, ILC2, however, BMAL1 was not required to establish cyclicity. Collectively, these data demonstrate that BMAL1 only partially influences ILC3 cyclicity and that other factors must be important in regulating this rhythm.

**Fig. 2.**
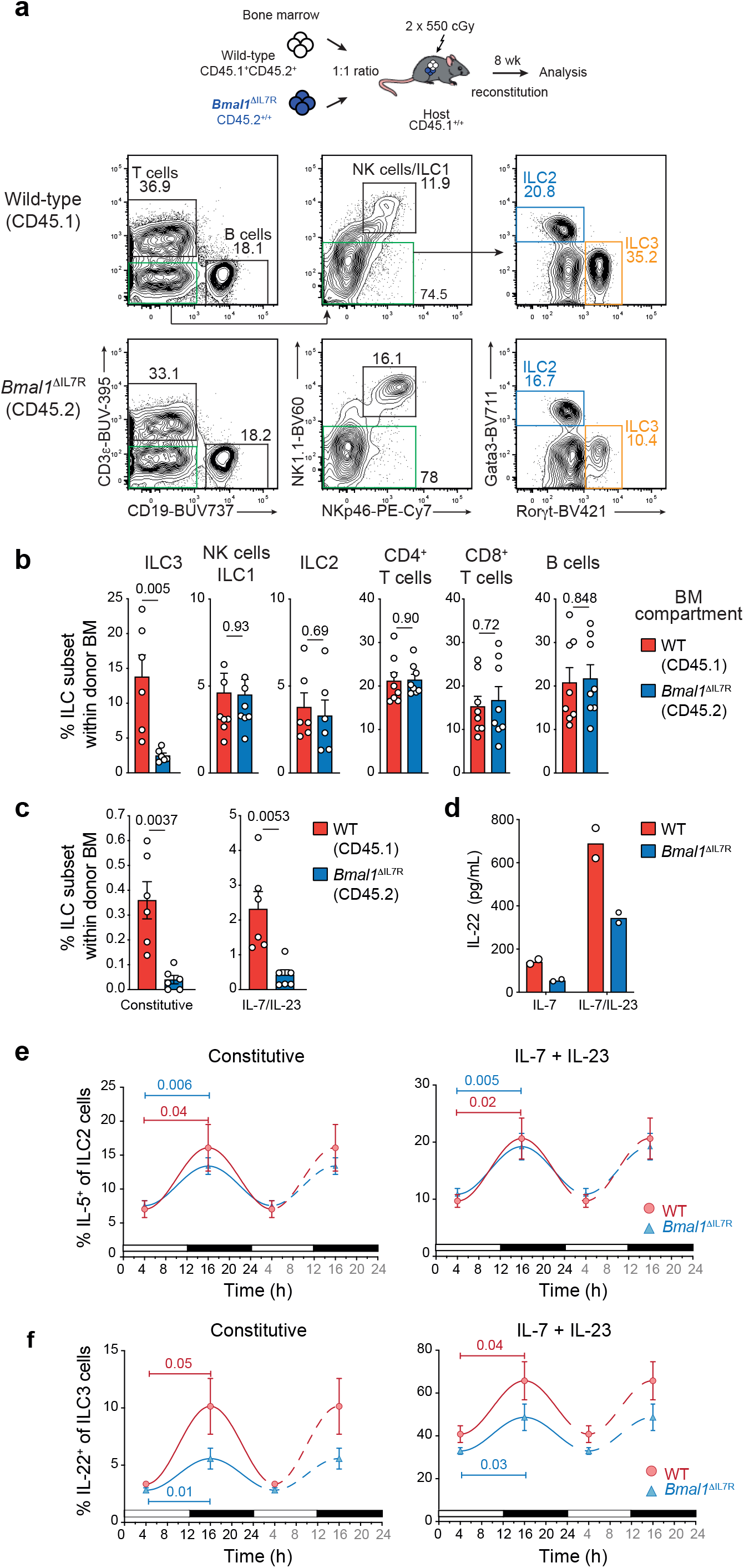
The molecular clock differentially but only partially regulates circadian expression of cytokine secretion. **a**-**c**, Mixed bone marrow chimeras were generated by reconstituting CD45.1^+/+^ lethally irradiated recipients with F1 (CD45.1 × CD45.2, CD45.1^+^CD45.2^+^) wild-type and CD45.2^+/+^ *Bmal1*^ΔIL-7R^ bone marrow. **a**, Representative contour plots of frequency of different lymphocyte subsets isolated in the small intestine of mixed bone marrow chimeras. Plots show live CD45.1^+^CD45.2^+^ cells for wild-type (F1) mice (upper panels) or CD45.2^+/+^ cells for *Bmal1*-deficient cells (lower panels). **b**, Frequency of different lymphocyte subsets derived from wild-type (red) and *Bmal1*-deficient (blue) donor bone marrow isolated from the small intestine of chimeric mice. **c**, Frequency of wild-type (CD45.1^+^CD45.2^+^, red) and *Bmal1*-deficient (CD45.2^+/+^, blue) IL-22-producing ILC3 isolated from the small intestine of chimeric mice. **a**-**c** Data are representive of 2 independent experiments (**a**) and show the mean ± s.e.m. for one experiment (*n* = 6-8 mice/experiment) (**b**,**c**). **d**, 4 × 10^3^ ILC3 were purified from the small intestine of wild-type (red) or *Bmal1*^*ΔIL7R*^ (blue) mice and cultivated 16 hours in presence of recombinant IL-7 (10ng/mL) to determine constitutive IL-22 expression, or IL-7 together with IL-23 (10 ng/mL) to elicit maximal IL-22 cytokine expression. IL-22 production was determined in culture supernatants by ELISA. **e-f**, Frequency of IL-5 and IL-22-producing (**e**) lung ILC2, and (**f**) small intestinal ILC3 at T4 and T16 isolated from wild-type and *Bmal1*^*ΔIL7R*^ mice. Left panel shows the constitutive expression of cytokines; right panel shows the cytokine expression after cultivation with IL-7 (10 ng/mL) and IL-23 (10 ng/mL). **e**,**f** Data show the mean ± s.e.m. of one of two similar experiments (*n* = 4 mice for each genotype per time point per experiment).

### Caloric intake is synchronized with ILC functional rhythmicity

Given that genomic deletion of *Bmal1* in ILC3 did not fully ablate rhythmic IL-22 cytokine production, we hypothesised that the cyclic behaviour of this subset in the gut may a depend on a combination of intrinsic and extrinsic factors. Because pathways involved in cellular metabolism were differentially regulated in our RNA-seq (Fig. 1c), we first mapped food intake across the light-dark phases of the 24 h period by tracking food consumption at different times (Fig. 3a). This revealed that the rate of food intake by mice was rhythmic and the pattern was similar to that observed for cytokine expression (Fig. 1b). Maximal food intake occurred during the dark phase while little food was consumed during the light phase. To understand whether food intake influenced ILC cyclic behaviour, food was provided either only during the dark (night food, high food intake) or the light (day food, low food intake) phase for seven days to adjust to this time-restricted feeding regime and establish new cellular cycles. The response of ILC2 and ILC3 were analysed at T4 after this period (Fig. 3b). For both intestinal ILC2 and ILC3, it was revealed that the frequency of ILC constitutively producing cytokines was increased when food was available during the dark phase (Fig. 3c, d). Interestingly, these differences were not observed in the lungs or the mesenteric LN, suggesting that the local regulation of ILCs in the intestine was directly dependent on food intake. We observed no differences in cytokine production in T cells (data not shown). To test whether the microbiota could be responsible for these differences, mice were treated with antibiotics for two weeks. Mice were given food only during the night, or only during the day, together with water containing antibiotics. Control mice received *ad libitum* access to food and water without antibiotics (Fig. 3e, blue bar). This quenched microbial signals and showed that antibiotic treatment during the active phase did not alter IL-22 production by ILC3 (Fig. 3e, red hatched blue bar). Both groups of mice that could consume food during the active phase showed an increased frequency of IL-22-producing cells compared with antibiotic treated mice fed only during the day (Fig. 3d, red hatched white bar) consistent with our earlier findings. Because all mice were analysed at T4, we wondered whether completely limiting access to food for a short period during the day would be sufficient to reduce cytokine expression in ILCs. To address this question we removed food for 16 hours prior to our analyses at T4 (Fig. 3f,g). This revealed that under time-restricted feeding, ILC2 and ILC3 isolated from the small intestine of fasted mice both showed reduced IL-5 (Fig. 3f) and IL-22 expression constitutively and following stimulation with IL-23, respectively, compared with fed mice, while ILCs from the mesenteric LN showed no changes (Fig. 3g). This effect was also observed in small intestinal ILC3 from fed and fasted animals indicating that the capacity of these cells to constitutively produce IL-22, and in response to IL-23, was imprinted *in vivo* (Fig. 3h). Combined, these results suggest that ILC3 responsiveness is highly tissue-specific and plays an important role in coordinating signals stimulated by food intake which overlies the intrinsic foundational circadian molecular machinery.

**Fig. 3.**
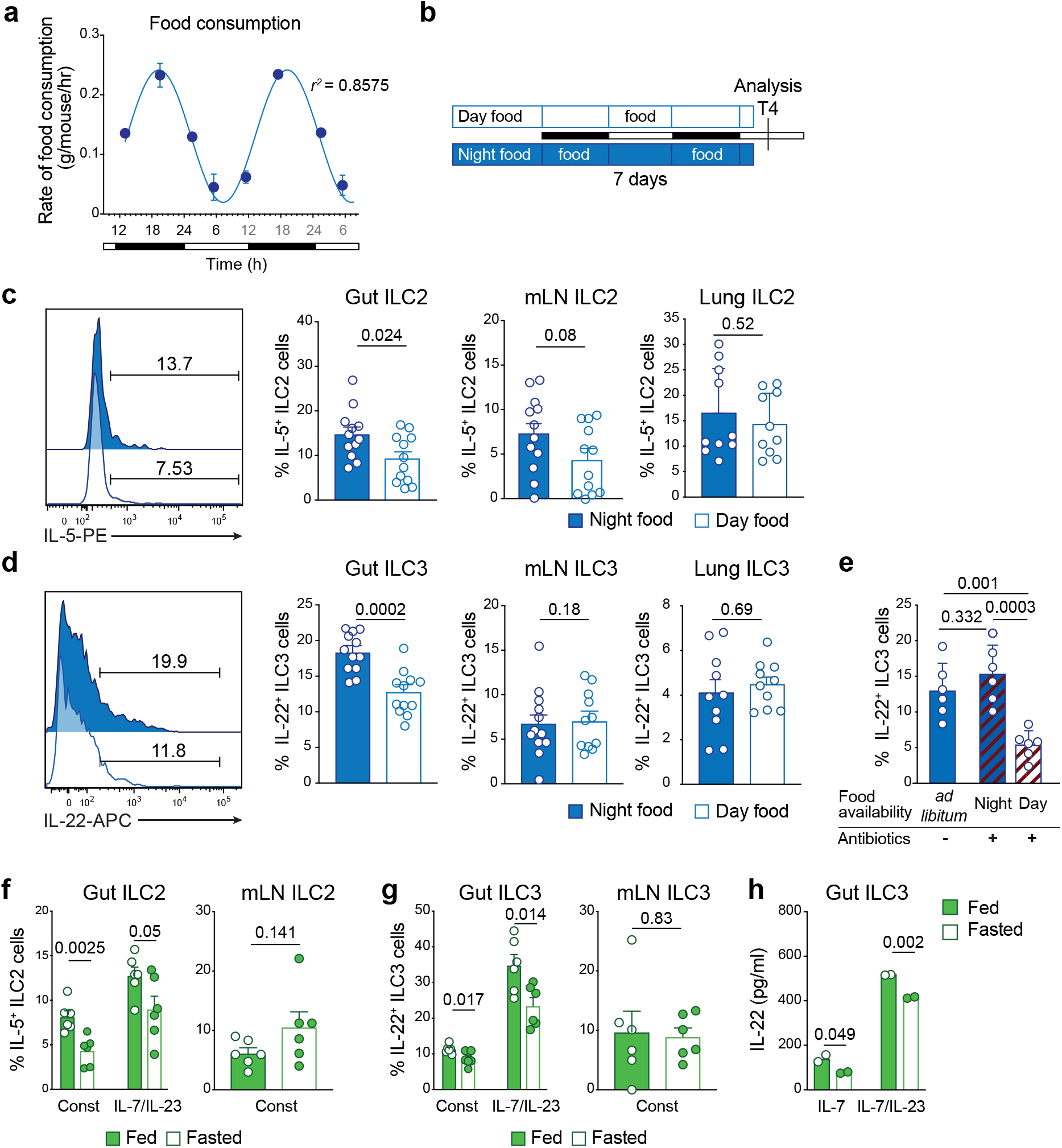
Food intake regulates the cytokine secretion of ILC2 and ILC3 in the small intestine. **a**, Food consumption of mice was determined at the indicated time points. Data show the mean ± s.e.m for one of two similar experiments (*n* = 12 mice per experiment). **b**, Schematic representation of the time-restricted feeding regime used in **c** and **d**. Mice were given food only during the day (day food) or the night (night food) for seven days. All mice were analysed at T4 on day 8. **c,d**, Analyses of IL-5-expressing ILC2 (**c**) and IL-22-expressing ILC3 (**d**) from the small intestine, mesenteric LN and lung of mice fed only during the day (white) or only during the night (blue) as determined by flow cytometric analyses of intracellular cytokine. Representiative flow cytometric histograms of cytokine expression (left panels) and the frequency of constitutively producing IL-5^+^ or IL-22^+^ ILC2 and ILC3, respectively (right panels), showing individual mice (circles) together with the mean ± s.e.m. pooled from two independent experiments (gut and mLN: *n* = 6, 6 mice for each condition/experiment; lung: *n* = 4, 6 mice for each condition/experiment). **e**, Mice were treated with combination antibiotics (ampicillin, streptomycin and colistin) for two weeks before analysis of constitutive IL-22 production by intracellular staining. Data show the mean ± s.e.m (*n*=6 mice per experimental condition). **f**,**g**, Expression of IL-5-expressing ILC2 (**f**) and IL-22-expressing ILC3 (**g**) from the small intestine and mesenteric LN from mice fed (green) or fasted for 16 hours (white). **f**,**g** Data show the mean ± s.e.m. of (*n* = 6 mice per group per condition) and show one of two similar experiments. **h**, Gut ILC3 (7.5 × 10^3^) were purified from fed (green) or fasted for 14 h (white) and cultivated 16 hours in presence of IL-7 (10 ng/mL) alone (constitutive expression) and IL-7 and IL-23 (10 ng/mL). IL-22 secretion was determined in culture supernatants by ELISA. Data show the response of cells pooled from 2 independent experiments with 3 mice per well for each condition in each experiment.

### Intestinal ILC3 differentially express the intestinal neuropeptide receptor VIPR2

To elucidate the potential mechanisms that might control the tissue-specific regulatory program observed in small intestinal ILCs, single cell RNA-seq was performed (Fig. 4a-c) on ILCs isolated from mice at T4. Analyses of data using t-Distributed Stochastic Neighbour Embedding (t-SNE)^38^ allowed us to visualize eight distinct clusters which included ILC1 and NK cells (cluster 7), ILC2 (cluster 4) and four different subsets of ILC3 (Clusters 1, 2, 3, 6) (Fig. 4a). This revealed that ILC3 in cluster 6 expressed the intestinal neuroendocrine receptor *Vipr2* (also known as *VPAC2*). This cluster mainly contained ILC3 expressing *Ccr6* and *Cd4* but were negative for *Ncr1* and *Tbx21* (Fig. 4b,c). While other ILC3 clusters constitutively expressed cytokines such as *Csf2* and *Il2* (cluster 1 and 2), the VIPR2^+^ ILC3 cluster mainly expressed *Il22* together with some *Il17* (Fig. 4c). In the colon, we observed only one cluster of ILC3 that constitutively expressed *Il22.* These ILC3 co-expressed *Ccr6*, *Cd4* and some cells expressed *Vipr2* (cluster 4, Supplementary Fig. 5). VIPR2 responds to VIP, a neuropeptide that is centrally involved in controlling cellular rhythms^39–41^ and is highly expressed in intestinal neurons to coordinate feeding responses^42^. More recently, VIP has been reported to drive IL-5 cytokine production in ILC2 for immune protection^28^. Thus, we hypothesised that expression of the receptor VIPR2 in ILC3 may act to coordinate the consumption of food intake with immune responsiveness of innate cells. Initially, we examined the localisation of VIP^+^ neurons in the small intestine by confocal and light sheet imaging of whole gut sections. Neurons were identified as β3-tubulin^+^ and co-stained with the neuronal mediator VIP that activates VIPR (Fig. 4d,e). This showed an extensive network of neurons (stained for β3-tubulin) extending from the the base of the intestinal crypts into the villus within the submucosa and adjacent to the lacteals. However, no VIP could be detected in the intestinal muscularis mucosae (Fig. 4d,e, Supplementary Movie 1). Staining of ILC3 (Rorγt^+^CD3ε^−^) revealed that intestinal neurons, VIP and ILC3 are closely connected anatomically positioning them for direct interations mediated by a feeding signal network (Fig. 4f). To determine whether food intake is synchronized with VIP expression, we next examined small intestinal sections for co-staining for intestinal neurons and VIP (Fig. 4g). This revealed that small intestinal tissue from fed mice showed co-localization of staining for neurons and VIP (shown in yellow/orange, Fig. 4g, left panel) in contrast to the situation in fasted mice (Fig. 4g, middle panel) where less VIP staining could be observed amongst neurons. Thus, fasting mice exhibited reduced expression of VIP in their small intestinal neurons while food intake induced secretion of VIP (Fig. 4g, right panel). This provides a direct link between VIP expression and ingestion of food as indicated by a strong correlation between feeding and VIP colocalization.

**Fig. 4.**
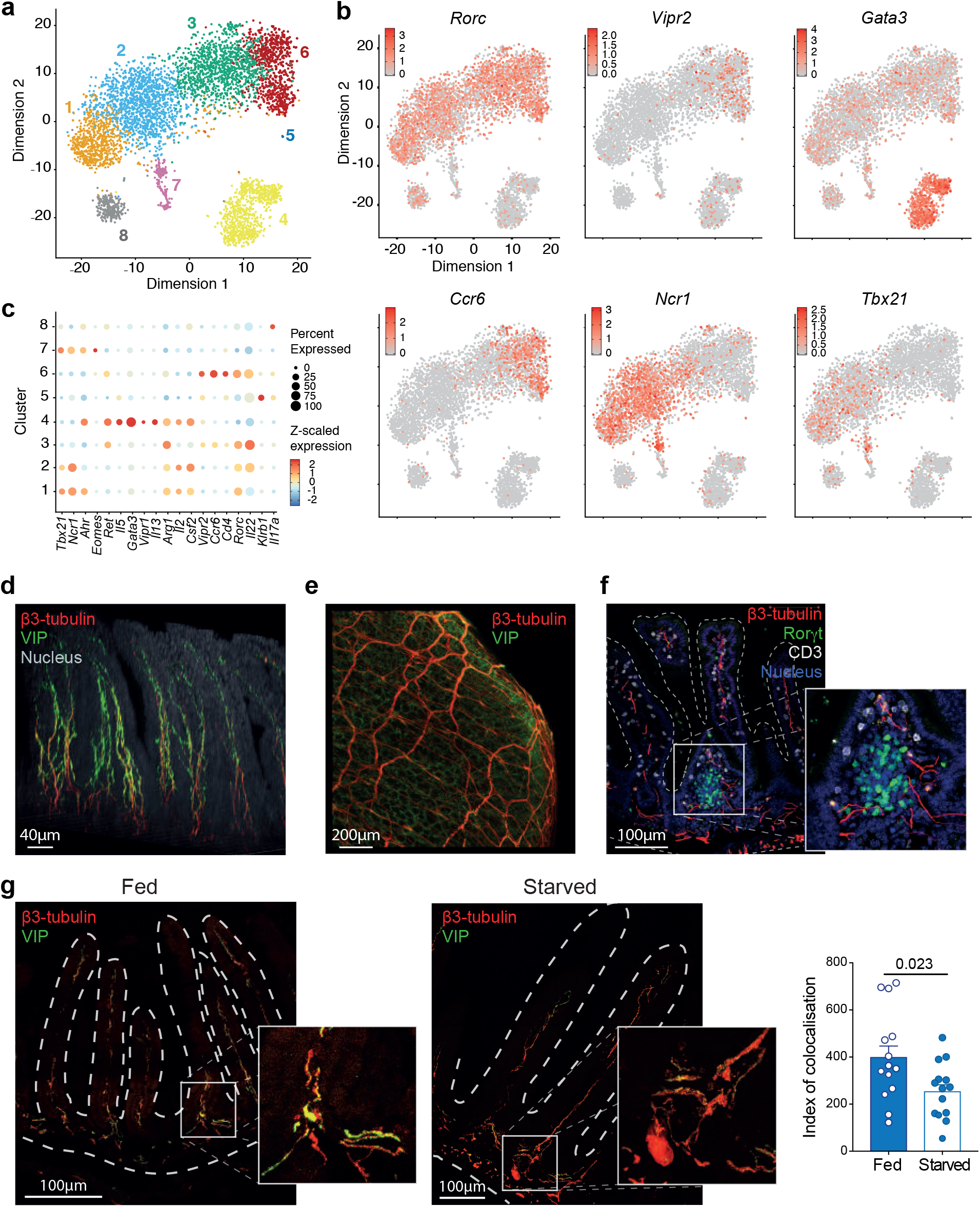
Intestinal ILC3 subsets express VIPR2 and are closely located to VIP-expressing neurons. **a**-**c**, ILC from the small intestine were sorted and single cells were sequenced using 10× Genomics. **a**, t-Distributed stochastic neighbour embedding (t-SNE) plots show 5,439 cells (dots) coloured by cluster. **b,** Level of expression of indicated genes within the t-SNE plots. **c**, Representative differentially expressed genes (x axis) by cluster (y axis). Dot size represents the fraction of cells within the cluster that express each gene. The colour intensity indicates the z-scaled expression of genes in cells within each cluster. **d**,**e**, Whole mount staining of the small intestine with β3-tubulin (red), VIP (green) and cell nuclei (blue). **d**, 3D reconstruction of confocal microscopy images showing the expression of VIP in neurons in the intestinal submucosa and the villi. **e**, Light sheet image showing VIP expression in the intestinal submucosa but not the intestinal muscularis mucosae. **f**, Confocal images of frozen sections of the small intestine from *Rorc(γt)*^*GFP/+*^ mice stained for neurons (β3-tubulin, red) ILC3 (GFP^+^ CD3ε^−^, green) and T cells (CD3ε^+^, white). Nuclei, blue. **g**, Colocalisation of VIP (green) in neurons (β3-tubulin, red) in the small intestine from fed and starved wild-type mice. Quantification of colocalization of β3-tubulin and VIP shows the the mean and s.e.m. of analyses of 14 different villi sampled from two mice per group (right panel).

### Feeding coordinates ILC3 IL-22 production and intestinal protection

To investigate the importance of the food intake induced VIP expression in detail, we assessed whether VIP exposure influences ILC3 activity. We stimulated intestinal cells with IL-7 and IL-23 in presence or absence of VIP. Dibutyryl cyclic-AMP (cAMP), a cell permeable cAMP analog that bypasses VIPR2 engagement to mimic direct cAMP stimulation and activate proten kinases^43^, was included as a positive control. Cells were obtained from mice that had been fasted for 16 hours to deplete VIP stores and hence their intrinsic capacity to stimulate ILC3. After 4.5 hours stimulation, intracellular expression of IL-22 was significantly increased in ILC3 cultivated in presence of synthetic VIP, or Dibutyryl cAMP (Fig. 5a). To determine whether VIP acted directly on ILC3, we purified small intestinal ILC3 from mice that had not been starved. These cells were cultivated in the presence of synthetic VIP, or dibutyryl cAMP for 16 hours (Fig. 5b). Interestingly, the addition of VIP to ILC3 cultures containing the survival factor IL-7 was sufficient to induce the secretion of IL-22, while the presence of IL-23 appeared to synergise with VIP to enhance IL-22 production (Fig. 5b). The influence of VIP, and dibutyryl cAMP, on IL-22 expression by ILC3 was dose dependent and occurred regardless of whether the induction of IL-22 was constitutive or provoked by the addition of IL-7 and IL-23 (Fig. 5c,d). To establish that VIP induction of IL-22 acted through the VIPR2 receptor itself, ILC3 were stimulated with the VIPR2 agonist BAY55-9837 (Fig. 5e). Strikingly, the response *via* VIP stimulation and in the presence of the VIPR2 agonist elicited similar IL-22 responsiveness in purified ILC3 while the VIPR2 antagonist PG99-465 largely blocked IL-22 secretion (Fig. 5f). Collectively, these findings connect the ingestion of food that drives VIP release in the small intestine with the coordinate enhancement of IL-22 secretion through VIPR2, a mechanism that would facilitate repair of tissue microabrasions of the intestinal epithelial that occur during feeding and exclude breach of the mucosal surface by microorganisms.

**Fig. 5.**
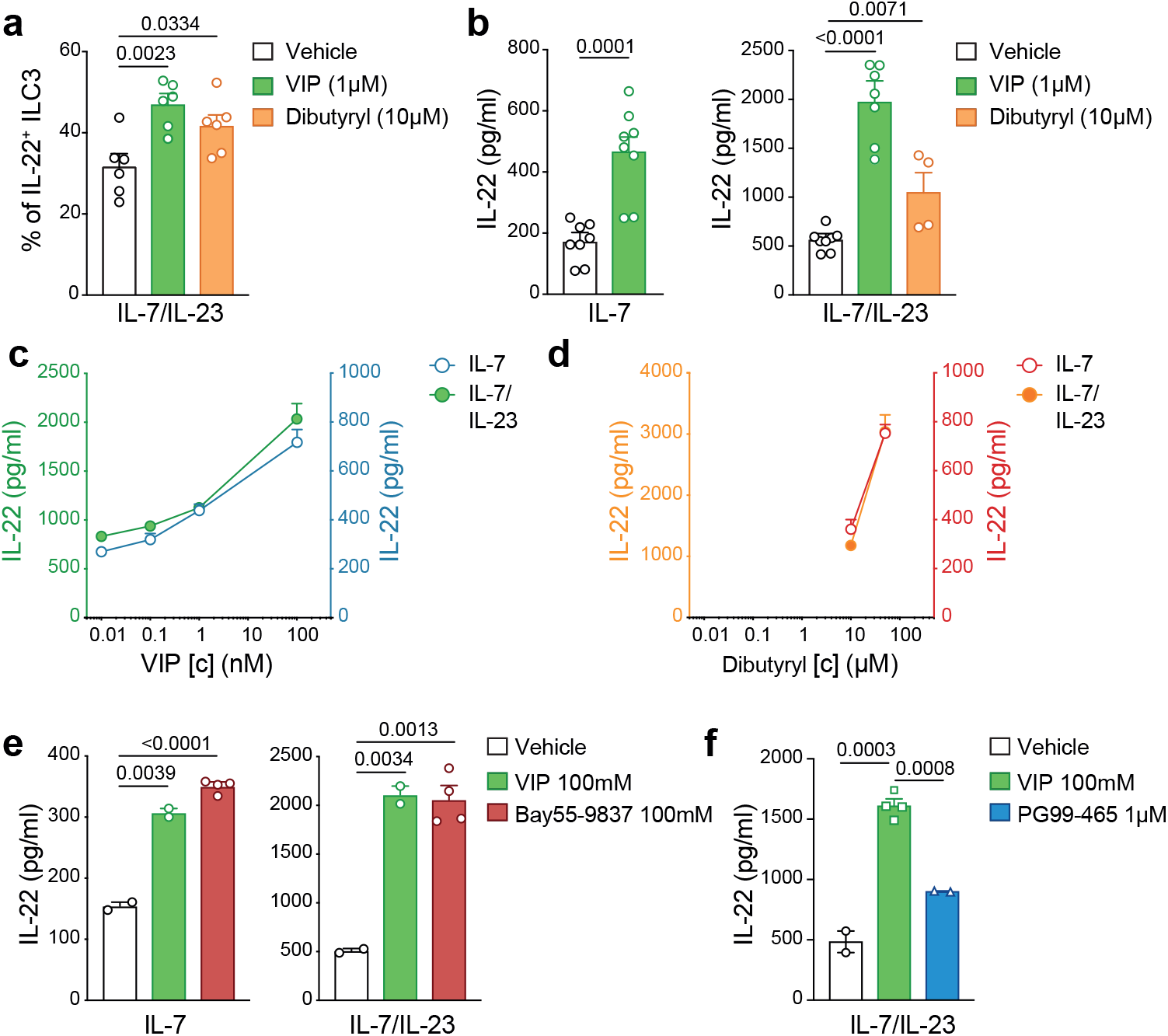
VIP directly regulates IL-22 production by ILC3. **a**, Intracellular expression of IL-22 detected in ILC3 after four hours of incubation with IL-7 and IL-23 (10 ng/mL) in presence of vehicle (DMSO), VIP (1 μM) or Dibutyryl cyclic-AMP (10 μM). **b**, IL-22 secretion by purified ILC3 isolated from the small intestine and cultivated for 16 hours with IL-7 (10 ng/mL) or IL-7 and IL-23 (10 ng/mL) in presence of vehicle (DMSO) or VIP (1 μM). IL-22 was measured in the supernatant by ELISA. Data are a pool of three independent experiments (mean ± s.e.m) from a total of six experiments. **c**,**d**, Titration effect of VIP (**c**) and Dibutyryl cyclic-AMP (**d**) on the production of IL-22 by intestinal ILC3. Purified ILC3 were stimulated with IL-7 alone or IL-7 and IL-23 (10 ng/mL) and different concentrations of either VIP (**c**) or dibutyryl cyclic-AMP (**d**). (**e**) IL-22 production of ILC3 cultured with IL-7 (10 ng/mL) or IL-7 and IL-23 (10 ng/mL) in the presence of VIP (100mM) or VIPR2 agonist BAY55-9837 (100mM). **f,** IL-22 production of ILC3 cultured with IL-7 and IL-23 (10 ng/mL) in the presence of VIP (100mM) or VIP (100mM) with the VIPR2 antagonist PG99-465 (1μM) for 16 hours. **e**,**f**, Data are pooled from two independent experiments, each dot represents a pool of two mice (mean ± s.e.m).

To definitively show that induction of IL-22 in ILC3 is mediated *via* VIP induction *in vivo*, mice were injected intraperitoneally with VIP, or PBS (control), 2 hours before analysis of the small intestinal ILC2 and ILC3 subsets (Fig. 6a). ILC3 exposed to VIP *in vivo* showed enhanced IL-22 production, but this treatment did not alter the induction of IL-17 secretion in these cells, or IL-5 secretion by in ILC2 (Fig. 6a). Mice deficient for VIPR2 (*Vipr2*^*−/−*^) showed a significant reduction in the proportion of ILC3 able to constitutively express IL-22 (Fig. 6b). To formally prove that VIP enhanced IL-22 production in ILC3 *via* VIPR2 signalling we purified ILC3 from wild-type and *Vipr2*^*−/−*^ and incubated the cells with IL-7 and IL-23 in presence or absence of VIP. After 16 hours of culture, we observed a strong upregulation of IL-22 secretion in wild-type cultures, while *Vipr2*^*−/−*^ ILC3 failed to respond to VIP (Fig. 6c). This failure to respond was not attributable to fewer ILC3 that could occur if VIPR2 was required for the development or maintenance of this subset as *Vipr2*^*−/−*^ mice had similar numbers of intestinal ILC compared with wild-type mice (Supplementary Fig. 6). Thus the VIP:VIPR2 axis is required for optimal timing of the induction of IL-22 secretion by ILC3.

**Fig. 6.**
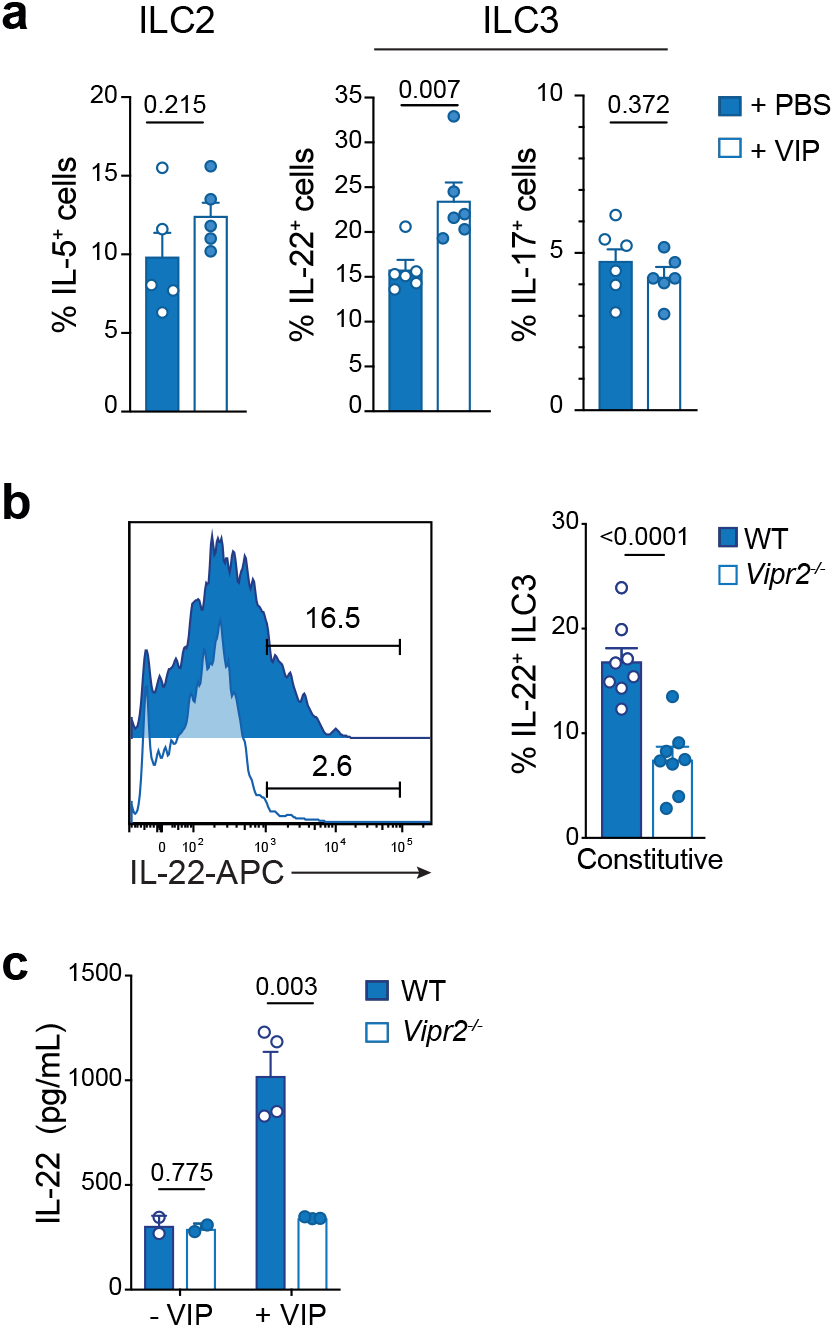
VIPR2 signalling regulates IL-22 secretion ILC3 *in vivo*. **a**, Mice were injected i.p. with 2 nM VIP or PBS 2h (T2) before sacrifice (T4). Constitutive expression of IL-5 by ILC2, and IL-22 and IL-17 by ILC3 from the small intestine was determined by flow cytometry. Data show the mean ± s.e.m. of one of two similar experiments (*n* = 5-6 mice per time point per experiment). **b**, Intracellular expression of ILC3 from the small intestine of wild-type or *Vipr2*^*−/−*^ mice after 4 hours of incubation with IL-7. Data show the mean ± s.e.m. of one of two similar experiments (*n* = 8 mice per time point per experiment). **c**, 4.5 × 10^3^ ILC3 purified from small intestine of wild-type or *Vipr2*^*−/−*^ were cultivated with IL-7 (10 ng/mL) alone or IL-7 and IL-23 (10 ng/mL) in presence of vehicle or VIP (1 μM) for 16 hours. IL-22 was measured in the supernatant by ELISA. Data represent a pool of 2 independent experiments and each dot represents a pool of two mice (mean ± s.e.m).

To understand whether VIP contributes to the physiological protection of the intestinal epithelium, we treated wild-type and *Vipr2*^*−/−*^ mice with low dose of dextran sodium sulfate DSS (1.5% w/v) for 5 days. While wild-type mice maintained their weight after 8 days of treatment, *Vipr2*^*−/−*^ animals started to lose weight after 5 days (Fig. 7a). *Vipr2*^*−/−*^ mice also showed more severe signs of colitis with shorter colon length (Fig. 7b) and reduced secretion of IL-22 by ILC3 (Fig. 7c). Histological analysis of the colon in *Vipr2*^*−/−*^ mice showed intestinal ulceration (Fig. 7d) together with significantly increased mucosal thickness (Fig. 7e), epithelial irregularity (Fig. 7f) and leukocyte infiltration (Fig. 7g) in *Vipr2*^*−/−*^ mice compared with wild-type mice. Thus, loss of VIPR2 signalling significantly impacted IL-22 production by ILC3 and the ability to protect against intestinal inflammation.

**Fig. 7.**
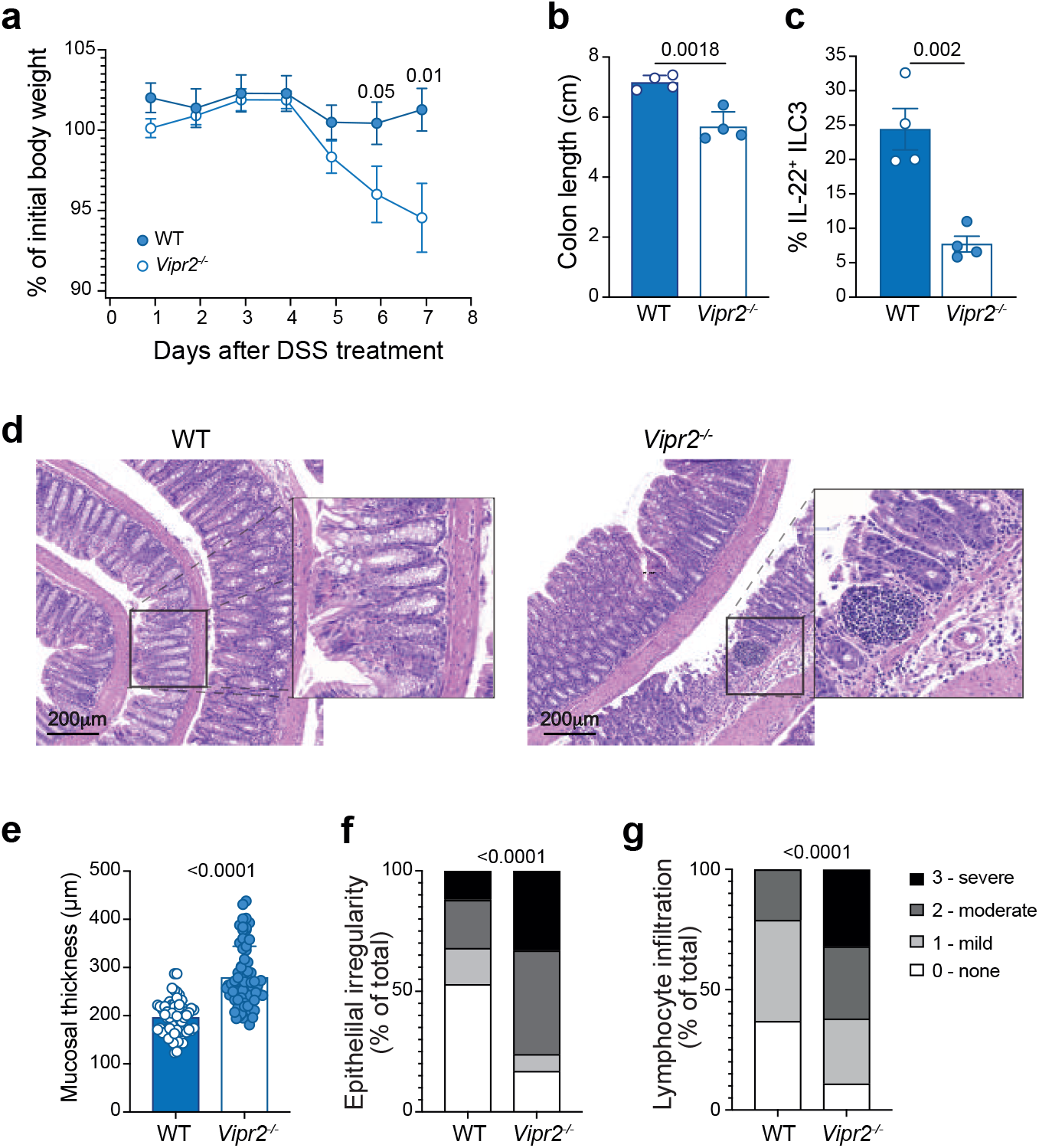
VIPR2 signalling is crucial to regulate gut integrity *in vivo*. **a-g,**Wildtype and *Vipr2*^*−/−*^ were given water containing 1.5% DSS (w/v) for 5 days followed by 3 days with regular water and analysed on day eight. **a**, Weight loss is shown as the mean ± s.e.m of percent initial weight of WT and *Vipr2*^*−/−*^ mice. **b**, Comparison of the colon length of wildtype and *Vipr2*^*−/−*^ treated with low dose DSS. **c**, Frequency of IL-22 positive ILC3 isolated from the small intestine of DSS-treated mice and cultivated 4.5 hours in presence of Golgistop. Individual responses (filled circles) together with the mean ± s.e.m for each genotype. **d**, Representative H&E stained sections of the colon from DSS-treated WT and *Vipr2*^*−/−*^ mice. Scale bar, 200 μm. **e-g**, Quantitative analysis of crypt height (**e**), epithelium irregularity (**f**) and inflammatory infiltrate (**g**) of the and width in WT and *Vipr2*^*−/−*^ mice. Epithelial length was measured in distal part of the colon. Overall epithelial irregularity and inflammatory infiltrate was scored from 6 sections over 2 different slides per mouse (0=normal, 1=minor, 2=moderate, 3=severe). Mucosal thickness was measured across 12 randomly selected sites points per mouse. Data show mean ± s.e.m (*n* = 4 mice/genotype) of one of two independent experiments with similar results.

## Discussion

ILC3 are an essential component of the intestinal barrier protection system. The cyclic regulation of IL-22 in the intestinal mucosa was dependent on food-induced neuronal VIP-VIPR2 ILC3 signaling to maintain immune repair after inflammation. While transcription factors associated with circadian clock machinery intrinsically imprint a rhythmic cycle in innate cells, we show that local cues such as feeding are also important and are regulated independently of clock machinery. This latter feedback loop is essential to coordinate the physiological fluctuations of the intestinal tract with immune protection.

Previous studies have shown that neuroimmune hubs composed of lymphocyte subsets, metabolic signals and neuronal coregulators play a key role in orchestrating context dependent immune protection^28, 31, 44–47^. In particular, ILC2 have been shown to be responsive to VIP through expression of VIPR2^28^. This pathway coordinates eosinophil recruitment and accumulation in the lung in a manner that parallels post-prandial induction of IL-5 secretion from intestinal ILC2 after feeding. Thus, we proposed that VIP-VIPR2 signalling predominates in ILC3. We found that IL-5 production by intestinal ILC2 was not intrinsically regulated by central clock machinery. In contrast, ILC3 were coregulated by intrinsic signals from BMAL1, and extrinsic signals from feeding, which itself was cyclicly regulated. As only part of the cell rhymicity program was governed by BMAL1, genetic ablation enabled us to uncouple the intrinsic pathway from extrinsic regulation of ILC3 activation revealing that ILC3-IL-22 functionality was tied to feeding patterns which would potentially also influence cellular metabolism and signalling.

As ILCs are thought to be almost exclusively lodged within tissues and positioned at barrier surfaces with little migratory capacity or ability to exert influence beyond the resident tissue^48, 49^, this constraint would likely significantly limit their capacity to exert broader protective functions without additional mechanisms driving the responsiveness of ILC with intrinsic and environmental stimuli. How then different mucosal barriers are protected by different ILC subsets has not been clear. Recently, it has been discovered that ILC2 could be mobilized from the bone marrow and traffick to the lung in response to helminth infection, a capacity regulated by sphingosine 1-phosphate (S1P)^50^ and dependent on the cyclic regulator BMAL1^51^. ILC2 are also regulated by the VIP-VIPR2 circuit to drive accumulation of key lymphocytes in the lung through the systemic release of IL-5^28^. Thus multiple, but distinct mechanisms, regulate ILC2 functionality allowing local and distant impacts on immune regulation. In contrast, we found that ILC3 responsiveness was highly tissue-specific and its IL-22 production was modulated by direct exposure to VIP from enteric neurons. Collectively, these data reveal that the gut ILC3-neuronal hub forms a very tightly regulated tissue-specific circuit executed exclusively by local cues. By deleting VIPR2 signalling, we demonstrated the critical requirement for the VIP-VIPR2 signalling pathway in eliciting ILC3-dependent IL-22 cytokine production to control inflammatory intestinal disease. This is consistent with the effector role of IL-22 in protecting against the development of inflammatory bowel disease and chronic colitis in both humans^52, 53^ and mouse models^10, 54, 55^. Thus, although the cellular source of IL-22 has been known for some time, it has not been understood the exact signals that allow for upregulation, or indeed downregulation which is equally important in limiting the immunopathology associated with excess IL-22.

Importantly, these data reveal a previously uncharacterised mechanism for coordinating how intestinal responses are matched to the stimulatory burden of the gut and orchestrate induction of homeostatic tissue repair pathways for mucosal protection. Furthermore, it highlights the differential contributions and mechanistic strategies utilized by ILC2 and ILC3 to finely tune local and distant ILC effector responses to orchestrate mucosal barrier protection, segregating homeostatic from proinflammatory functions of these cells. Our present findings shed new light on how VIPR2 expressed by ILC3 confer immediate tissue protection through coordinated neuroimmune sensory responses establishing their unique role in coordinating food-induced VIP secretion with IL-22-driven cytokine protection.

## Methods

### Mice

*Bmal1*^*fl/fl*^ (B6.129S4(Cg)-*Arntl*^*tm1Weit*^/J)^56^, *Il7r*^*Cre* 57^, *Vipr2*^*−/−*^ (B6.129P2-*Vipr2*^*tm1Ajh*^/J)^58^, *Rorc(γt)*^*+/GFP* 59^ and *Rag2*^*−/−*^*γc*^*−/− 60*^ mice were maintained on a C57BL/6 (Ly5.2) background (all originally derived from the Jackson Laboratory) and have been described previously. *Bmal1*^*fl/fl*^ mice were crossed to the *Il7r*^*Cre*^ strain to generate the *Bmal1*^*fl/fl*^*Il7r*^*Cre*^ line that lacks BMAL1 in all lymphocytes. C57BL/6, B6.SJL-*Ptprc*^*a*^*Pep3*^*b*^/BoyJ (Ly5.1^+/+^, CD45.1^+/+^) and CD45.1^+^CD45.2^+^(F1) mice were bred and maintained in-house. Mice were housed under a 12 h:12 h light-dark cycle in temperature controlled conditions with food and water *ad libitum* unless otherwise indicated. For time-restricted feeding experiments, food was removed for a period of 12 hours either during the day, or the night with water available *ad libitum*. For some experiments mice were fasted overnight for a period of 16 hours. When not stipulated mice were analysed at time (T) 4 hours after the onset of light and food access was not restricted. Mice were bred and maintained in specific pathogen free conditions at the Walter and Eliza Hall Institute of Medical Research. All mice were female and were used at 7-12 weeks old. Animals were handled according to the guidelines of the Australian code for the care and use of animals of the National Health and Medical Research Council of Australia. Experimental procedures were approved by the Animal Ethics Committee of the Walter and Eliza Hall Institute of Medical Research.

### Isolation of lymphoid cells

Single cell suspensions of lymphocytes were isolated from the small intestine and colon following incubation for 40 min at 37°C in Ca^2+^- and Mg^2+^-free Hanks media containing 2% FCS and 1 mM EDTA with gentle shaking to remove intestinal epithelial cells^5, 61^. The supernatant was discarded and tissues were then incubated, with gentle shaking, for 40 min with 1mg/mL (w/v) Collagenase type IV (Worthington), 200 μg/mL DNase I (Roche), 4 U/mL Dispase (Sigma) in RPMI-1640 + 2% (v/v) heat inactivated foetal calf serum (FCS) for 45 min at 37°C. Lungs were cut into small fragments and digested for 45 min at 37°C with 1 mg/mL Collagenase IV and 200 μg/mL DNase I. Preparations were filtered and mononuclear cells were isolated by centrifugation on a 40%-80% Percoll gradient. Lymphocytes were recovered from the interface and washed twice. Single cell suspensions were generated from mesenteric LN by gently dissociating tissues using 70μm filters. Cells were resuspended in PBS supplemented with 1mM EDTA.

### Flow Cytometry

Single-cell suspensions were stained with the following antibodies: TCRβ (H57-597) from BioLegend; CD19 (ID3), CD3ε (145-2C11), ICOS (C398.4A), NKp46 (29A1.4), NK1.1 (PK136), CD117 (2B8), CD127 (IL-7R, A7R34), Sca-1 (D7), and ST2 (RMST2-2) from eBioscience; and CD45.1 (A20) CD45.2 (104), CD90.2 (30-H12), and CD49a (Ha31/8) from BD Biosciences. Intracellular staining was performed using the Transcription Factor Staining Buffer Set (eBioscience) and antibodies against GATA-3 (TWAJ), Rorγt (AFKJS-9, BD Biosciences) and EOMES (Dan11mag) (eBioscience) and cytokines IL-5 (TRFK5, eBioscience), IL-13 (eBio13A, eBioscience), IL-17A (eBio17B7, eBioscience) and IL-22 (IL22JOP, eBioscience). Cells were analysed using a FACS Canto II or Fortessa X20 (BD Biosciences) and FlowJo software was used for analysis. Flow cytometric sorting was performed with a FACSAria III (BD Biosciences).

### ILC3 isolation and *in vitro* stimulation

Cells isolated by digestion from the small intestinal tissue were stained for 30 min with optimally-titred antibodies against CD90.2, NKp46, NK1.1, CD117, IL-7Rα, KLRG1, B220, CD19, CD3, CD11b and CD45. ILC3 were identified as CD45^+^CD90^+^CD117^+^IL-7Rα^+^NKp46^+/−^ and negative for KLRG1, B220, CD19, CD3, CD11b and NK1.1. Stained cells were washed and sorted using an FACS ARIA III (BD Biosciences). Purified cells were cultivated in complete RPMI-1640 containing 10% heat inactivated FCS, 1mM L-glutamine, 100U/mL penicillin, 100 μg/mL streptomycin and 50μM β-mercaptoethanol. The cytokine expression of ILCs was determined using the following conditions: cells were stimulated for 4.5 hours in the presence of 1ng/mL phorbol-12-myristate-13-acetate (PMA), 1μg/mL ionomycin and 10μg/mL Brefeldin A and GolgiStop™ (BD Biosciences). To test the constitutive secretion of cytokines, ILCs were incubated 4.5h in presence of 10 μg/mL Brefeldin A. For longer term cultures, media was supplemented with 10 ng/mL recombinant human IL-7 (R&D Systems)(a survival factor), with or without 10ng/mL recombinant murine IL-23 (R&D Systems). To test the effect of VIP on ILC3, cells were incubated with different concentrations of VIP (human, rat, mouse, rabbit, canine, porcine) from Tocris, Bay 55-9837 (Tocris), PG99-465 [Myristoyl-(Lys12.27·28) VIP-Gly-Gly-Thr (free acid) trifluoroacetate salt](Bachem) or dibutyryl-cAMP (Tocris). All regeants were resuspended in sterile water, except dibutyryl-cAMP that was resuspended in DMSO. To measure IL-22, purified ILC3 were cultured for 16 hours in the above conditions and the supernatant analysed using the IL-22 ELISA Kit (Thermo Fisher Scientific).

### *In vivo* activation with exogenous VIP

For all *in vivo* experiments with VIP treatment, an intraperitoneal *(i.p*.) administered dose of 2 nmol VIP in PBS per mouse or PBS alone was injected two hours before tissue harvest.

#### DSS-induced colitis

Dextran sodium sulfate (DSS) (molecular mass 36,000 – 50,000 Da; MP Biomedicals) was added into drinking water 1.5% (w/v) for five days followed by three days of regular water. Mice were analysed on day eight. Mice were monitored daily for weight loss, stool consistency and hematochezia.

### RNA sequencing and analyses

ILC2 and ILC3 cells were purified by flow cytometric sorting (FACS Aria, BD Biosciences) at four different time points (T4, T10, T16, T23) from female wild-type mice. One to two biological replicates were generated for each subset from small intestine. RNA was isolated using a QIAGEN RNeasy Micro kit. Libraries were generated using the Illumina Truseq RNA sample preparation kit following the manufacturer’s instructions and subjected to transcriptome 85 bp single end sequencing on an Illumina Next-Seq instrument. Reads were aligned to the GRCm38/mm10 build of the *Mus musculus* genome using the Rsubread aligner^62, 63^. Uniquely mapped reads were retained and genewise counts were obtained using featureCounts^64^ and NCBI RefSeq annotation. Differential expression analyses were then undertaken using the edgeR^65^ and limma^66^ software packages. Lowly expressed genes were filtering using edgeR’s filterByExpr function with default settings. Additionally genes without current annotation, ribosomal RNAs, non-protein coding immunoglobulins, predicted/unknown/pseudo genes, and those not on known chromosomes were also removed. Compositional differences between libraries were normalised using the trimmed mean of log expression ratios (TMM) method^67^. To remove unwanted variation from the data and adjust for differences between the ILC2 and ILC3 samples, the RUVIII alogorithm from the ruv software package^68^ was applied with k=4 to the log_2_ counts per million (CPM) transformed data. A single harmonic sinusoidal model was then fit to the corrected data and cyclic genes were identified using robust empirical bayes moderated t-statistics with a trended prior variance^69^. P-values were adjusted using the Benjamini and Hochberg method to control the FDR below 5%. Analysis of the Reactome database was carried out using limma’s kegga function. The heatmap was generated using the CRAN package pheatmap version 1.0.12.

### Single cell RNA sequencing and analysis

For single cell RNA sequencing (scRNA-seq), ILC were sorted as lin^−^(CD3ε^−^TCRβ^−^CD19^−^ B220^−^Gr1^−^CD11b^−^) CD45^+^IL-7R^+^CD90^+^ cells into ice cold 0.5% BSA in PBS and processed through the Chromium Single Cell 3′ v2 Library Kit (10× Genomics) according to the manufacturer’s protocol. Each channel was loaded with 20,000 cells from each tissue, yielding 5439–7388 single cells for analysis from small intestine and colon. The cells were then partitioned into gel beads in emulsion in the instrument, where cell lysis and barcoded reverse transcription of RNA occurred, followed by amplification, shearing, and 5′-adaptor and sample index attachment. Libraries were sequenced on an Illumina NextSeq platform. Reads from each sample were separately processed using 10× Genomics’ Cell Ranger software (v3.0.2). Cellranger mkfastq was used to demultiplex the Illumina sequencer’s base for each flowcell directory into FASTQ files. Next, ‘cellranger count’ was used to generate single cell gene count matrices against 10× Genomics’ pre-built mm10 reference genome and transcriptome (v3.0.0). All subsequent analysis was performed in R (v3.6.0)^70^ and Bioconductor (v3.9)^71^ and simpleSingleCell workflow^72^. The DropletUtils package (v1.4.0) was used to create a *SingleCellExperiment* for each sample from the Cell Ranger output directories and to identify empty droplets^73^. Quality control on the cells was performed using the scater package (v1.12.2)^74^ with two rounds of outlier detection and removal. Cells were declared as outliers and removed if they satisfied any of the following criteria: Read counts more than three median absolute deviations (MADs) below the median (log10 scale); Number of genes detected more than three MADs below the median (log10 scale); Percentage of reads coming from the mitochondria more than three MADs above the median. Detection of doublet libraries was performed using a 2-stage strategy similar to that of Pijuan-Sala *et al.*^75^. In total, this identified 262 putative doublets (211 from the Fed Colon and 51 from the Fed SI) which were removed from subsequent analysis. We then performed principal component analysis (PCA) across all samples and used the mutual nearest neighbours corrected values, we applied t-stochastic neighbourhood embedding (t-SNE) using the Rtsne package (v0.13)(https://cran.r-project.org/web/packages/Rtsne). To identify cluster-specific marker genes, we performed Welch *t*-tests on the log-expression values for every gene and between every pair of clusters.

### Histology and confocal imaging

For histological analyses, tissues were fixed in 10% neutral buffered formalin and processed for wax embedding and routine hematoxylin and eosin (H&E) staining. Tissues were analyzed with an Aperio ImageScope v11.2.0.780 software and features were blindly scored for the presence of mucosal ulceration, mucosal height, epithelial irregularity and lymphocyte infiltration using the grading system: 0=no changes, 1=mild changes, 2= moderate changes and 3=severe changes.

For confocal imaging analysis, intestinal tissues were gently flushed with PBS then cut longitudinally and formed into a ‘Swiss roll’ for visualization of the full length of the tissue. After fixation of 4% paraformaldehyde for two hours, the Swiss rolls were dehydrated overnight in 30% sucrose prior to embedding in OCT, frozen and stored in 80°C. Frozen tissues were sliced using a cryostat to 20 μm thick sections and fixed in cold acetone for 10 min. Tissues were washed with PBS and blocked with PBS containing 3% BSA, 0.03% Triton X-100 for two hours at room temperature. Tissue sections were then rinsed in staining buffer (1% BSA, 0.03% Triton X-100 in PBS) and incubated with anti-β-tubulin III Alexa 594 (TUBB3, Biolegend), anti-VIP Alexa 647 (H-6, Santa-Cruz) and anti-CD3ε Pacific blue (OKT3, eBioscience), Hoechst 33342 (Thermofisher) and SYTO^®^ 24 (Invitrogen). Tissues were stained overnight at 4°C with antibodies diluted in PBS containing 1% BSA, 0.03% Triton X-100 (Sigma). Tissue sections were then washed three times then mounted with ProLong Gold mounting medium (Invitrogen). Images were acquired on a Zeiss LSM880 NLO confocal microscope.

For whole mount staining, small intestinal tissues were cleaned and cut in small sections (~ 0.5 cm). Sections were fixed in 4% PFA for two hours at room temperature. Afterwards the tissue was washed in PBS and blocked with PBS containing 3 % BSA and 0.3% Triton X-100 for 12 hours. The tissue was stained for 48 hours at 4 °C with the following antibodies: anti-β-tubulin III Alexa 594, anti-VIP Alexa 647 and anti-CD3ε Pacific blue or Hoechst 33342. Subsequently gut samples were cleared using Ce3D reagent^76^. The tissue was then embedded in low-melting temperature agarose (Bio-Rad) in a 1 mL syringe to be imaged with a Zeiss Z.1 light sheet (Zeiss) microscope or mounted on a microscope slide and captured under a Zeiss LSM 880 NLO confocal microscope equipped with Zen software (Zeiss). Surface reconstruction was performed by using Imaris software (Version 9.3) (Bitplane). Data analyses was *via* a custom macro in FIJI^77^.

### Oral antibiotic treatment

Mice were given unrestricted access to drinking water containing a mix of ampicillin (1 mg/mL), streptomycin (5 mg/mL) and colistin (1 mg/mL) (Sigma) for two weeks^78–80^. Gut decontamination was evaluated by macroscopic evaluation of caecal hypertrophy.

### Statistical analysis

All statistical analyses were performed with GraphPad Prism software (Version 7.0 or 8.0, Graphpad Software). An unpaired two-tailed Student’s *t* test was used for pairwise comparisons. An unpaired and paired Student’s t-test, nonparametric Mann-Whitney test or Log-rank Mantel-Cox test was used for other comparisons. All experiments were performed at least two times with similar results obtained for each time. For visualising circadian patterns, we performed COSINOR fitting of circadian curves using the curve-fitting module of Graphpad Prism with the equation Y = baseline + amplitude × cos (Frequency X + Phaseshift), where Baseline = average of Ymax and Ymin; Amplitude = 0.5 x (Ymax − Ymin), Frequency=0.2618 (2*π/24) and Phaseshift= value of X at Ymax.

### Data availability

Single cell RNAseq profiling data that support the findings of this study have been deposited in the Gene Expression Omnibus (GEO) repository with the accession code GSEXXX.

## Acknowledgements

We wish to thank Mary Camilleri, Ann Lin, Shelby Cree, Carolina Alvarado and Tracy Putoczki for expert technical advice and support. We thank Stephen Wilcox for performing sequencing and Stephen Nutt for critical reading of the manuscript. Financial support for this work was provided by National Health and Medical Research Council (NHMRC) of Australia grants (APP1165443, 1122277, and 1054925 G.T.B. and C.S.), Cure Cancer and Cancer Australia (APP1163990 N.J.), The Rebecca L. Cooper Foundation Medical Research Foundation (G.T.B.), fellowships from the NHMRC (APP1135898 G.T.B., APP1123000 C.S., APP1154970 G.K.S.). This study was made possible through Victorian State Government Operational Infrastructure Support and Australian Government NHMRC Independent Research Institute Infrastructure Support scheme.

The authors declare no competing interests.

## Author contributions

C.S., K.L., A.L.G., P.H., N.J., J.T., V.C.W. and R.D.S. performed experiments and data analyses. G.K.S., A.L.G., P.H. and M.R. oversaw bioinformatic analyses. V.C.W., K.R. and L.W. oversaw imaging analyses. G.T.B. and C.S. directed the studies and wrote the paper.

## Author Information

The authors declare no competing financial interests.

