## Supplementary data for "Vasoactive intestinal peptide confers anticipatory mucosal immunity by regulating ILC3 activity"

**Supplementary Fig. 1-6**
**Supplementary Movie 1**

**Supplementary Figures**

**Supplementary Fig. 1. Cytokine secretion of ILCs at T4 and T16**

**a**,**b**, Frequency of cytokine-producing NK cells, ILC1, ILC2 and ILC3 isolated from the lungs (**a**) and mesenteric LN (**b**) at T4 and T16. IFN-γ, TNF-α, IL-5, IL-13, IL-17 and IL-22 production was determined by intracellular cytokine staining in the indicated populations. Individual responses together with mean ± s.e.m. are shown. **a**,**b**, *n* = 7-8 mice at each time point. Data is shown for one of four to six similar experiments.

**Supplementary Fig. 2. Cytokine secretion of T lymphocytes at T4 and T16**

**a-c,** Frequency of indicated cytokine produced by $CD4^+$ T cells and $Th17^+$ cells ($CD4^+Ror\gamma t^+$) in the small intestine (**a**), lungs (**b**) and mesenteric LN (**c**) from naïve C57BL/6 mice at T4 and T16. IFN-γ, TNF-α, IL-17 and IL-22 production was determined by intracellular cytokine staining. Individual responses together with mean ± s.e.m. are shown. **a-c,** *n* = 7-8 mice at each time point. Data show one representative of four to six similar experiments.

**Supplementary Fig. 3. Enhanced cytokine expression in ILC subsets at T4 and T16.**

Geometric mean fluorescence intensity of intracellular IL-5 and IL-22 cytokine production from ILC2 and ILC3, respectively, isolated from the small intestine of naïve C57BL/6 mice at T4 and T16. Shown are individual responses together with mean ± s.e.m. (*n* = 4 mice per time point) for one of four similar experiments.

**Supplementary Fig. 4. Purification of small intestinal ILC3 from food-restricted mice.**

**a**,**b**, ILC3 were isolated from the small intestine of food-restricted mice by high-speed flow cytometric sorting of $lin^-(CD3\varepsilon^-TCR\beta^-CD19^-B220^-Gr1^-CD11b^-)CD45^+IL\text{-}7R^+CD90^+c\text{-}kit^+$ cells using a BD FACSARIA III (BD Biosciences). **a**, Representative dot plots show the cell surface markers and gating strategy used to discriminate ILC3 from other ILC subsets. **b**, Representative analyses of ILC3 purity after cell sorting.

**Supplementary Fig. 5. Single cell RNA-sequencing of colon ILC3.**

**a-c,** ILC from colon were sorted and single cells were sequenced using 10× Genomics. **a**, *t*-Distributed stochastic neighbour embedding (t-SNE) plots show 7,388 cells (dots) coloured by cluster. **b,** Level of expression of indicated genes within the t-SNE plots. **c**, Representative differentially expressed genes (x axis) by cluster (y axis). Dot size represents the fraction of cells within the cluster that express each gene. The colour intensity indicates the z-scaled expression of genes in cells within each cluster.

**Supplementary Fig. 6. Total number of small intestinal lymphocytes in *Vipr2*$^{-/-}$ mice.**
Enumeration of ILC1, ILC2, ILC3, B cells, CD8$^{+}$ T cells, CD4$^{+}$ T cells and Th17$^{+}$ T cells isolated from the small intestine of naïve wild-type and *Vipr2*$^{-/-}$ mice. Data show the mean ± s.e.m. of one of two similar experiments ($n$ = 8 mice per time point per experiment).

**Supplementary Movie 1**.
Whole mount staining of the small intestine of C57BL/6 mice with antibodies against β3-tubulin (red) and VIP (green). Z stack imaging was reconstructed in three dimensions using the Imaris software. Imaging shows the intestinal nerve network (red) from the muscularis mucosae to the tip of the villi. VIP is shown in green in enteric nerves from the crypt to the tip of the villi.

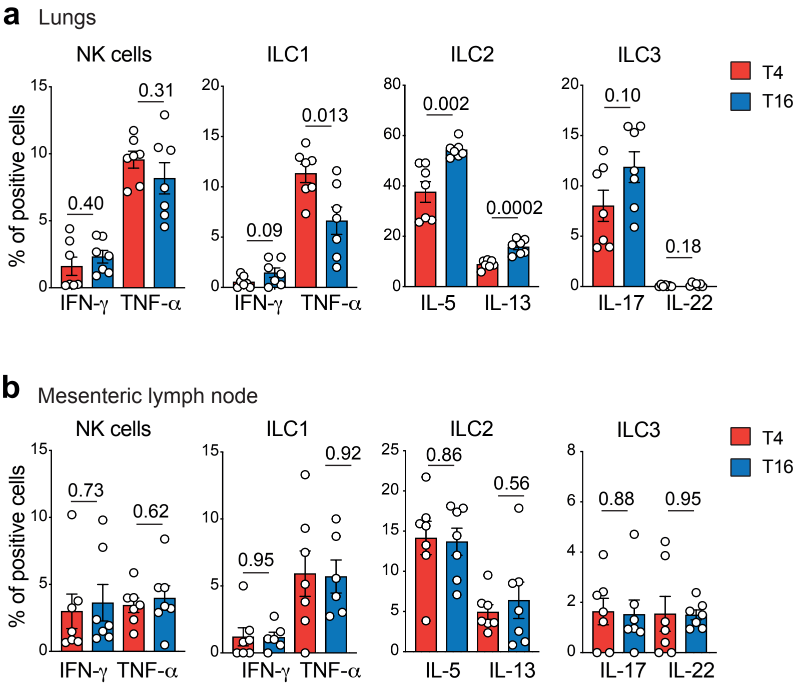


Supplementary Figure 1, Seillet *et al.*

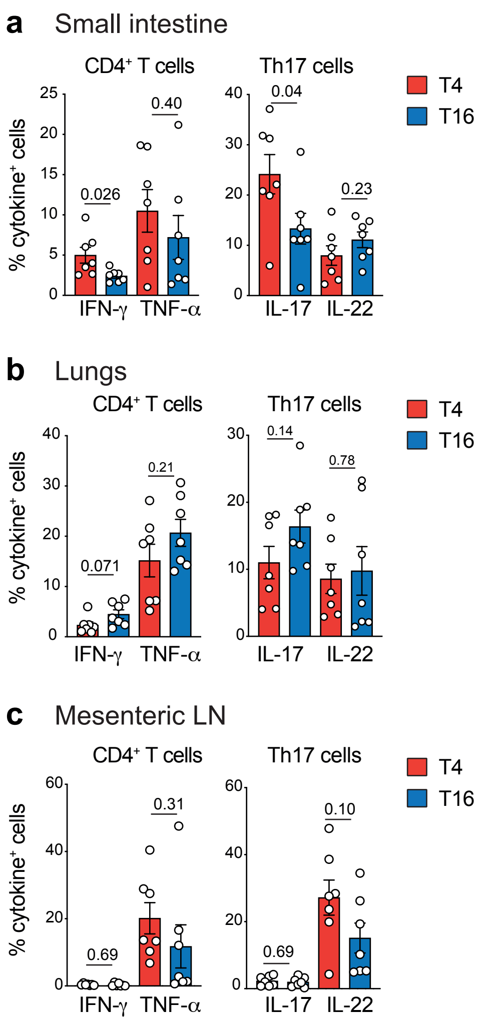


Supplementary Figure 2, Seillet *et al.*

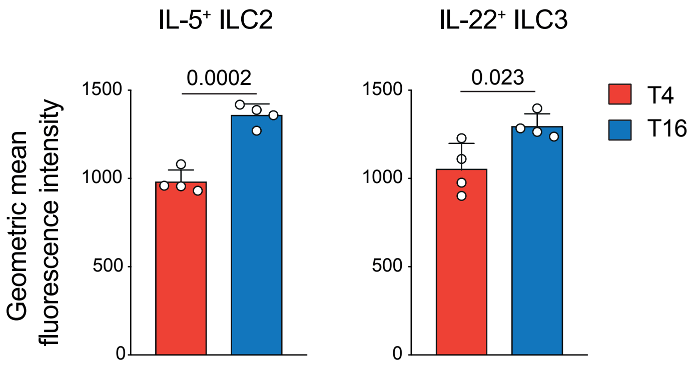


Supplementary Figure 3, Seillet *et al.*

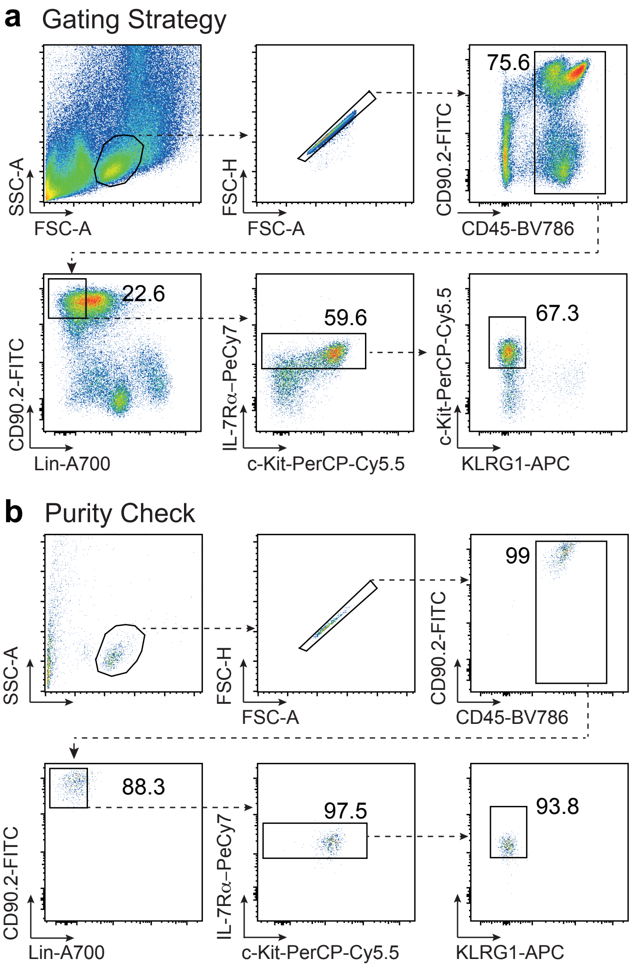


Supplementary Figure 4, Seillet *et al.*

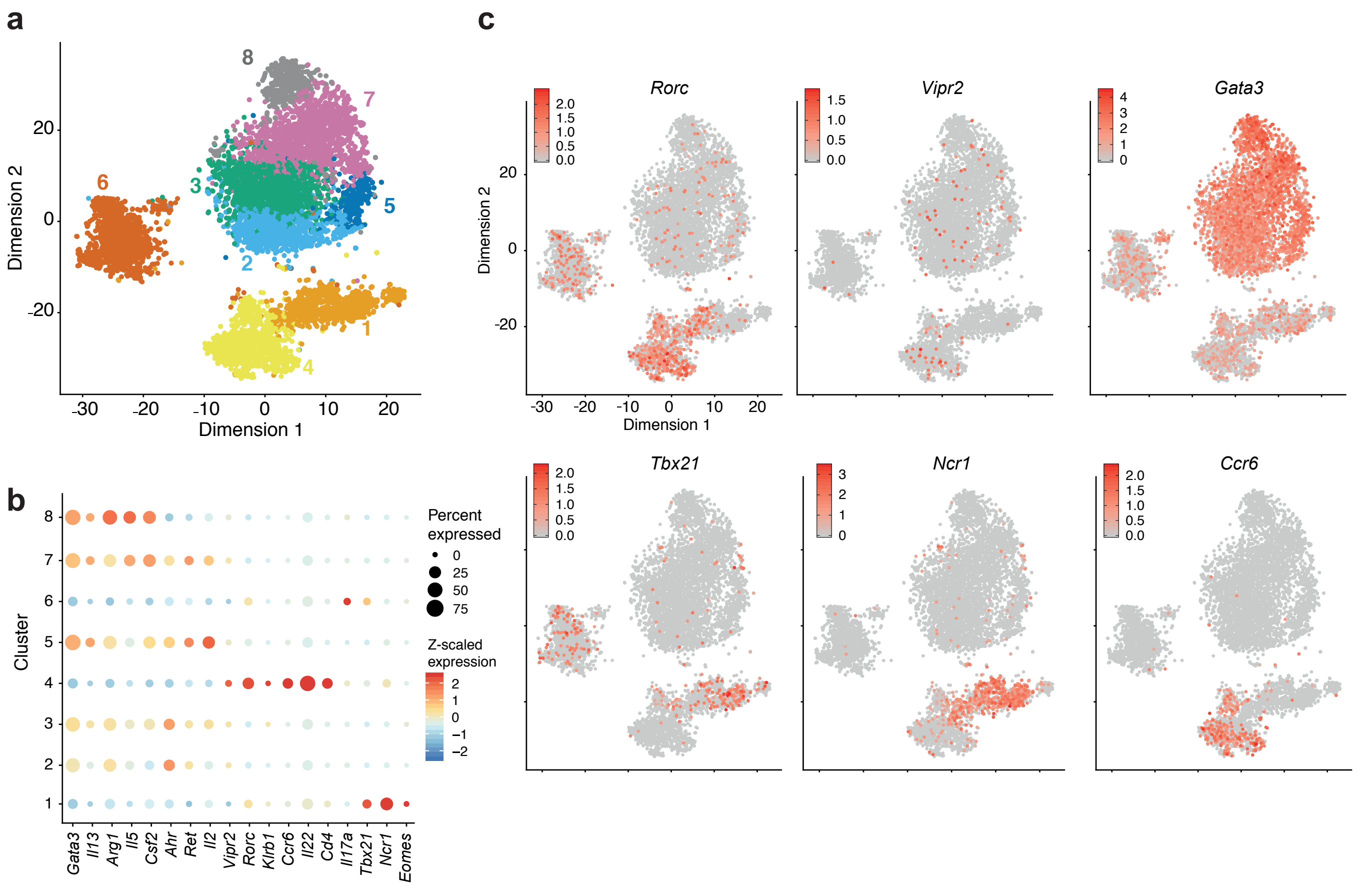


Supplementary Figure 5, Seillet *et al.*

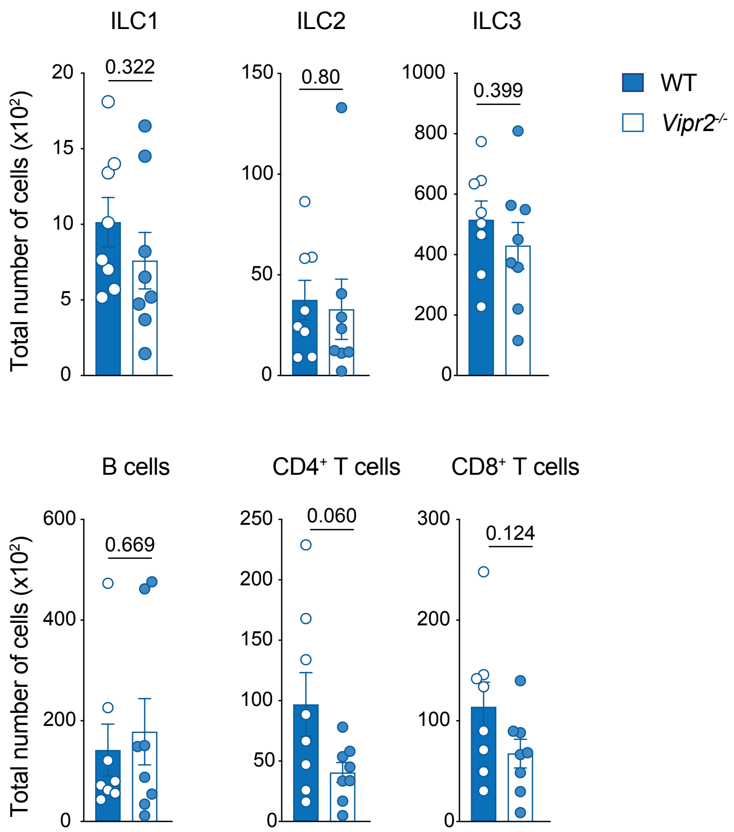


Supplementary Figure 6, Seillet *et al.*
